# Discovery of a broadly-neutralizing human antibody that can rescue mice challenged with neurotoxin-rich snake venoms

**DOI:** 10.1101/2022.06.17.496531

**Authors:** Line Ledsgaard, Jack Wade, Kim Boddum, Irina Oganesyan, Julian Harrison, Timothy P. Jenkins, Pedro Villar, Rachael A. Leah, Renato Zenobi, John McCafferty, Bruno Lomonte, José M. Gutiérrez, Andreas H. Laustsen, Aneesh Karatt-Vellatt

## Abstract

Snakebite envenoming continues to claim many lives across the globe, necessitating the development of improved therapies. To this end, human monoclonal antibodies may possess advantages over current plasma-derived antivenoms by offering superior safety and improved neutralization capacity. However, as new antivenom products may need to be polyvalent, *i.e*., target multiple different snake species, without dramatically increasing dose or cost of manufacture, such monoclonal antibodies need to be broadly-neutralizing. Here, we report the establishment of a pipeline for the discovery of high affinity broadly-neutralizing human monoclonal antibodies. We further demonstrate its utility by discovering an antibody that can prevent lethality induced by *N. kaouthia* whole venom at an unprecedented low molar ratio of one antibody per toxin, and which also prolongs survival of mice injected with *Dendroaspis polylepis* or *Ophiophagus hannah* whole venoms.

## INTRODUCTION

Each year, snakebite envenoming exacts a high death toll and leaves hundreds of thousands of other victims maimed for life^1^. Antivenoms based on polyclonal antibodies isolated from the plasma of immunized animals are currently the only specific treatment option against severe envenomings^2,3^. While these medicines are essential and life-saving, and will remain a cornerstone in snakebite therapy for years to come, an opportunity now exists to modernize treatment by exploiting the benefits of recombinant DNA and antibody technology^4^. Indeed, recombinant antibodies have already been generated against a variety of snake venom toxins^5–8^. Moreover, within this area of research, it has been demonstrated that monoclonal antibodies targeting snake venom toxins can be developed using various platforms, such as phage display technology^6^, an *in vitro* methodology that can be used to actively select for antibodies with high-affinity and cross-reactivity^7,8^. In addition, the use of human antibody libraries in combination with phage display technology allows for the discovery of fully human antibodies, that are likely to have high treatment tolerability in patients.

It has been speculated that monoclonal antibodies developed by these means could be used to formulate recombinant antivenoms that elicit fewer adverse reactions, are cost-competitive to existing therapy, and can be fine-tuned to have superior efficacy^9–13^. Phage display technology could be particularly valuable for discovering monoclonal antibodies against highly potent toxins with low immunogenicity that fail to elicit a strong antibody response in animals used for immunization^14,15^. This is the case for low molecular mass neurotoxins and cytotoxins of the three-finger toxin (3FTx) family, which are abundant in Elapidae venoms, such as cobra and mamba venoms^16–19^. However, antibodies derived directly from naïve libraries often lack sufficiently high affinity to enable toxin neutralization^8^. Affinity can be improved by further site-directed or random mutagenesis of the antibody paratopes, which can also lead to broadening of the neutralizing capacity of naïve antibodies^20^. However, besides mutation of the antibody binding regions, retaining the heavy chains and exploring alternative light chains, a technique known as light chain-shuffling, has shown significant promise as well^14,21^. Here, a phage display library is generated by paring a heavy or light chain from a specific antibody with a naïve repertoire of the partner chain and performing a new selection campaign^8^. Nevertheless, until now it remained unknown whether this technology could be used to generate antibodies that possess high affinity while simultaneously having a broad neutralization capacity, *i.e*., are able to neutralize several related toxins from the venoms of different snake species.

Previously, using a naïve human scFv-based phage display library^8^, we described the discovery and characterization of the human monoclonal antibody, 368_01_C05, against α-cobratoxin (P01391), a potent neurotoxin from the monocled cobra, *Naja kaouthia*. Notably, this antibody could prolong the survival of mice injected with lethal doses of α-cobratoxin, although it failed to prevent lethality^8^. As a follow-up development, in the present study we constructed light-chain-shuffled antibody libraries based on this clone with the aim of using a phage display-based cross-panning campaign to simultaneously improve the affinity and expand the neutralizing capacity of the antibody against α-neurotoxins from the venoms of several snake species. Cross-panning was carried out between α-cobratoxin^22^ and α-elapitoxin^23^, a neurotoxin from the venom of the black mamba, *Dendroaspis polylepis*^18^. These two α-neurotoxins share 70% sequence identity and both cause neuromuscular blockade by binding to the nicotinic acetylcholine receptor (nAChR) in muscle cells.

In this work, we “cross-panned” the chain-shuffled scFv library on these two toxins under stringent conditions to discover antibodies with improved affinity and cross-reactivity in comparison to the parent antibody. Using this strategy, we were able to generate an antibody which not only has improved affinity to α-cobratoxin, but also significantly broadened cross-neutralization capacity against other α-neurotoxins from the venoms of elapid snakes from the genera *Dendroaspis, Ophiophagus, Bungarus*, and *Naja*.

## RESULTS

### Affinity maturation, cross-panning selections, and scFv characterization

Human light-chain-shuffled scFv-based phage display libraries were created by paring the heavy chain of antibody 368_01_C05 with a naïve repertoire of human light chains. Then, phage display cross-panning selections using two toxins with 70% sequence identity, α-cobratoxin from *N. kaouthia* and α-elapitoxin from *D. polylepis*, were conducted according to the overview provided in Fig. 1A. Phage display selection outputs from the third round were subcloned into the pSANG10-3F expression vector, and 736 monoclonal scFvs were tested for their ability to bind to α-cobratoxin, α-elapitoxin, and streptavidin in both direct dissociation-enhanced lanthanide fluorescence immunoassays (DELFIAs) and expression-normalized capture (ENC) DELFIAs. From here, 203 scFvs (all displaying binding to at least one of the two toxins with negligible binding to streptavidin) were randomly selected for sequencing. Out of these, 67 scFvs were unique according to the sequence of their light chain CDR3 region, 2 of them having kappa light chains and the remaining 65 having lambda light chains. The top 62 clones, based on sequence diversity and binding behavior, were reformatted to the fully human IgG1 format. Following expression in HEK293 cells, ENC DELFIAs were run using the crude expression supernatant to rank the IgG binding to α-cobratoxin, α-elapitoxin, a venom fraction from *N. melanoleuca* (Nm8) containing a long neurotoxin homologous to OH-55 (Q53B58) and long neurotoxin 2 (P01388)^24^, as well as streptavidin. This revealed that more than half of the clones were cross-reactive against all three toxins/venom fractions, demonstrating significant improvement in both binding and cross-reactivity when compared to the parental antibody.

**Fig. 1.**
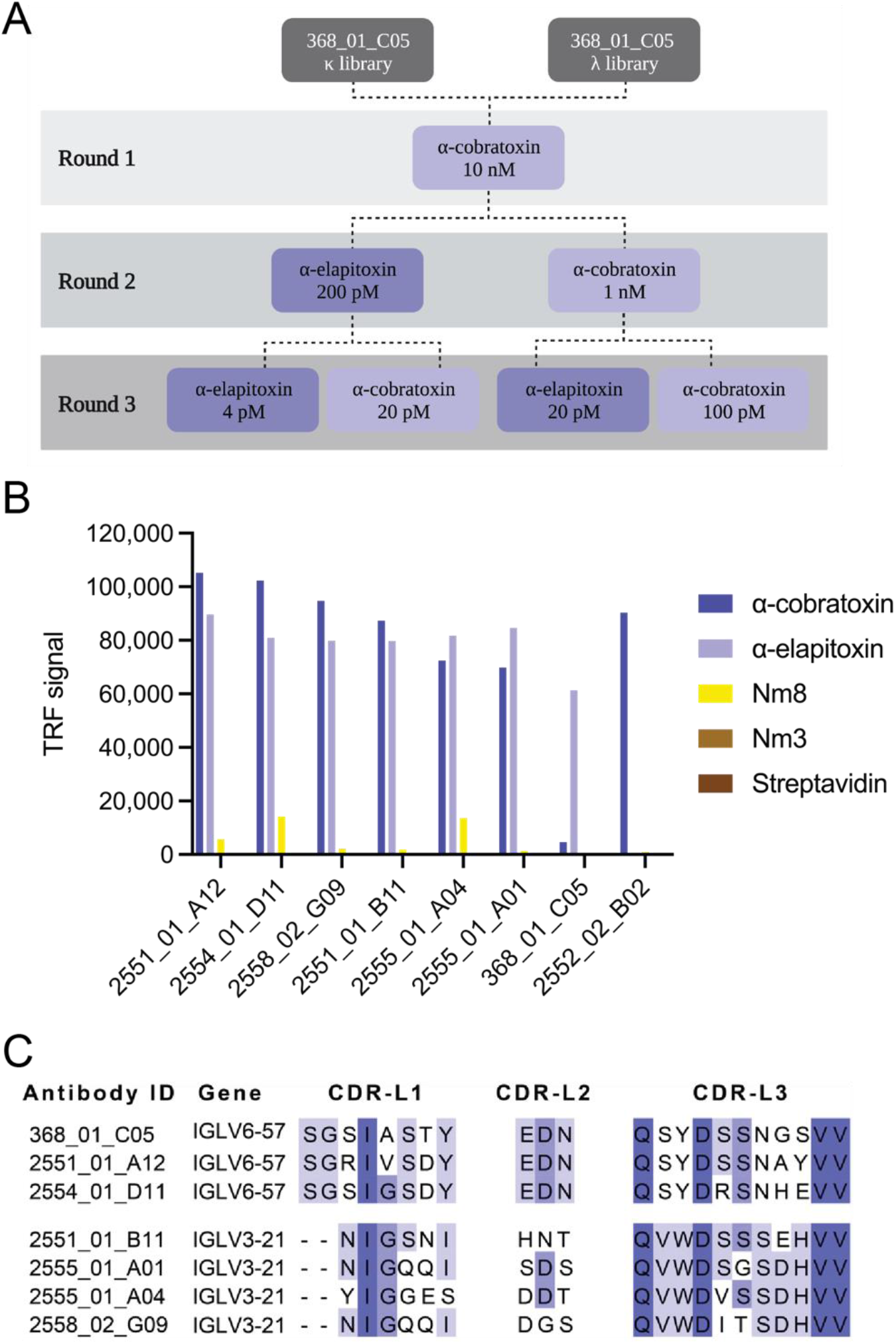
Cross-panning selection strategy as well as assay and sequence data for selected IgGs. A) Selection strategy illustrating how cross-panning was performed, including antigen concentrations. B) ENC DELFIA showing cross-reactivity of top six affinity matured IgGs (2551_01_A12, 2551_01_B11, 2554_01_D11, 2555_01_A01, 2555_01_A05, and 2558_02_G09) in comparison with parental IgG (368_01_C05) and clone 2552_02_B02 from a previously published study^8^. C) Comparison of CDR-L, CDR-L2, and CDR-L3 sequences for the top six chain-shuffled antibodies and the parental antibody.

To help guide the selection of lead candidates, the suitability of the 62 clones for future antibody development was investigated by characterizing biophysical properties that are indicative of their “developability” profiles. To this purpose, we analyzed the purity and non-specific column interaction pattern of all IgGs using size-exclusion chromatography (SEC). In addition an AC-SINS assay^25^ was employed to asses propensity for self-aggregation. For this analysis, we also included an IgG from a previous study (2552_02_B02)^8^, that had been reported to neutralize lethality induced by *N. kaouthia* whole venom *in vivo*, but had never been characterized for cross-reactivity to other long chain α-neurotoxins nor been analyzed for its “developability” properties. The SEC data (% monomeric content and relative elution volumes - a metric for assessing non-specific interaction with the SEC column), AC-SINS shifts, binding data, expression yields (full dataset see Table S1), as well as light chain germ-line diversity were used to select the top six antibody candidates for further characterization. These antibodies were named as follows: 2551_01_A12, 2551_01_B11, 2554_01_D11, 2555_01_A01, 2555_01_A04, and 2558_02_G09 (Fig. 1B and Table 1). Additionally, the data showed that the previously published IgG 2552_02_B02 had an extremely poor developability score, both judged by its late elution in the SEC analysis and its high shift in the AC-SINS assay. In fact, this antibody performed at a similarly poor level as the ‘poor developability’ control (bococizumab, AC-SINC shift of 33 nm) that was used for comparison, whereas all antibodies derived from the 368_01_C05 parental clone possessed developability profiles similar to the ‘good developability’ control antibody (Aliricumab, AC-SINS shift of 3 nm). In addition, 2552_02_B02 showed no cross-reactivity to any of the long chain α-neurotoxins it was tested against, clearly distinguishing its binding profile from the antibodies derived from the 368_01_C05 parent (Fig. 1B).

**Table 1.** **AC-SINS shift, SEC analysis results (% monomer content and relative elution volume), and transient expression yields** for the top six chain-shuffled IgGs (2551_01_A12, 2551_01_B11, 2554_01_D11, 2555_01_A01, 2555_01_A05, and 2558_02_G09) in comparison with the parental IgG (368_01_C05). IgG 2552_02_B02 from a previous study was also included.

| Antibody ID | AC-SINS | SEC analysis |  | Production |
| --- | --- | --- | --- | --- |
|  | Shift (nm) | Monomer (%) | Elution (mL) | Yield (mg/L) |
| 2551_01_A12 | 3 | 100.0 | 1.48 | 18.1 |
| 2554_01_D11 | 1 | 94.8 | 1.48 | 47.1 |
| 2558_02_G09 | 1 | 95.6 | 1.48 | 38.5 |
| 2551_01_B11 | 1 | 96.2 | 1.50 | 30.4 |
| 2555_01_A04 | 1 | 100.0 | 1.47 | 25.8 |
| 2555_01_A01 | 1 | 96.7 | 1.47 | 45.4 |
| 368_01_C05 | 0 | 97.5 | 1.45 | 30.0 |
| 2552_02_B02 | 32 | 100.0 | 1.82 | 22.6 |

Analysis of the antibody sequences revealed that the six affinity matured antibodies had light chains belonging to two different germlines, germline IGLV3-21 for 2551_01_B11, 2555_01_A01, 2555_01_A04, and 2558_02_G09 and germline IGLV6-57 for 2551_01_A12 and 2554_01_D11. The parental antibody had germline IGLV6-57, meaning that two of the six affinity matured antibodies had light chains belonging to the same germline as the parental antibody. From the comparison of the three light chain CDR regions of the antibodies presented in Fig. 1C, it could also be seen that for the two antibodies maintaining the parental germline, the CDR-L2 was identical to the parental, whereas the CDR-L1 and CDR-L3 had 2-3 amino acid differences. For the remaining four antibodies with different light chain germline, all VL CDR sequences were significantly different from the parent antibody sequence.

To evaluate if the light-chain-shuffling campaign generated antibodies with improved affinity, surface plasmon resonance (SPR) was used to determine the affinity of the top six antibodies as well as the parental antibody. To this purpose, all antibodies were reformatted to the monovalent Fab format to measure 1:1 binding kinetics of each antibody against both α-cobratoxin and α-elapitoxin (for SPR sensograms see Fig. S1). Data showed that all six antibodies displayed higher affinity to both toxins as compared to the parental antibody (Table 2). The largest improvement was observed for 2551_01_A12 and 2554_01_D11 (32 and 50-fold improvement of binding to α-cobratoxin and 13 and 8-fold improvement of binding to α-elapitoxin, respectively), providing both antibodies with low single-digit nanomolar affinities to both toxins. Thus, a significant improvement in both affinity and cross-reactivity was observed. Antibodies 2551_01_A12 and 2554_01_D11 were selected for further characterization based on affinity, cross-reactivity, expression yield, and developability data.

**Table 2.** Affinity measurements using Surface Plasmon Resonance (SPR). SPR was used to measure the affinity of the top six chain-shuffled antibodies (2551_01_A12, 2551_01_B11, 2554_01_D11, 2555_01_A01, 2555_01_A05, and 2558_02_G09) and the parental antibody (368_01_C05) in the Fab format to both α-cobratoxin and α-elapitoxin. The dissociation constants, on-rates, and off-rates are provided. For sensograms see Fig. S1.

| Antibody ID | α-cobratoxin | | | α-elapitoxin | | |
| --- | --- | --- | --- | --- | --- | --- |
|  | K<sub>D</sub> (nM) | k<sub>on</sub> (M·s) | k<sub>off</sub> (s<sup>-1</sup>) | K<sub>D</sub> (nM) | k<sub>on</sub> (M·s) | k<sub>off</sub> (s<sup>-1</sup>) |
| 2551_01_A12 | 2.79 | 1.80·10<sup>5</sup> | 5.02·10<sup>-4</sup> | 1.12 | 1.25·10<sup>5</sup> | 1.40·10<sup>-4</sup> |
| 2554_01_D11 | 1.78 | 1.28·10<sup>5</sup> | 2.29·10<sup>-4</sup> | 1.69 | 1.05·10<sup>5</sup> | 1.77·10<sup>-4</sup> |
| 2558_02_G09 | 2.77 | 2.14·10<sup>5</sup> | 5.94·10<sup>-4</sup> | 3.04 | 1.39·10<sup>5</sup> | 4.22·10<sup>-4</sup> |
| 2551_01_B11 | 4.27 | 1.17·10<sup>5</sup> | 5.00·10<sup>-4</sup> | 2.87 | 6.37·10<sup>5</sup> | 1.83·10<sup>-4</sup> |
| 2555_01_A04 | 7.46 | 2.22·10<sup>5</sup> | 1.65·10<sup>-3</sup> | 2.21 | 9.26·10<sup>5</sup> | 2.05·10<sup>-4</sup> |
| 2555_01_A01 | 8.41 | 2.02·10<sup>5</sup> | 1.70·10<sup>-3</sup> | 1.81 | 1.00·10<sup>5</sup> | 1.81·10<sup>-4</sup> |
| 368_01_C05 | 89.6 | 1.83·10<sup>4</sup> | 1.64·10<sup>-2</sup> | 14.3 | 6.47·10<sup>4</sup> | 9.25·10<sup>-4</sup> |

Because the binding profiles of the cross-reactive antibodies derived from antibody 368_01_C05 were significantly different from antibody 2552_02_B02, SPR was used to determine if the antibodies bound the same or overlapping epitopes on α-cobratoxin (see Fig. S2). Using 2554_01_D11 as a representative of the cross-reactive antibodies, this study revealed that neither of the two antibodies 2552_02_B02 or 2554_01_D11 could bind α-cobratoxin if the other antibody was already bound to the toxin, suggesting that the antibodies recognized the same or overlapping epitopes.

### Native mass spectrometry reveals cross-reactivity to several toxins from elapid snakes of three different genera

To further explore the cross-reactivity of the discovered antibodies, IgG 2554_01_D11 was tested for its binding to toxins in five whole venoms including *N. kaouthia*, *N. melanoleuca*, *N. naja*, *Ophiophagus hannah*, and *D. polylepsis*. These venoms from African (*D. polylepsis* and *N. melanoleuca*) and Asian (*N. kaouthia*, *N. naja* and *O. hannah*) snakes all possess a relatively high content of long chain α-neurotoxins, ranging from 13.2% for *D. polylepsis*^18^ to 55% for *N. kaouthia*^17^, except *N. naja* that has been reported to have a long chain α-neurotoxin-content of about 2-5%^26^. For this purpose, native mass spectrometry (MS), was used to investigate the interactions between the antibody and toxins from the four snake venoms.

Prior to native mass spectrometry analysis, the venoms and the IgG were fractionated using SEC (Fig. S3). The IgG was mixed with each of the SEC-generated toxin fractions, before analysis using native MS to determine binding (Fig. S3). This analysis revealed that 2554_01_D11 only bound toxins of masses in the range expected for the group of three-finger toxins (3FTx) to which all α-neurotoxins belong. To identify the toxins, the toxin:antibody complexes were isolated using MS/MS and subjected to collisional dissociation to eject the toxins from the antibody, allowing their intact mass to be determined. The primary dissociation product from these experiments were proteins of masses between 7,800 and 8,200 Da, corresponding to typical masses of long chain α-neurotoxins (Fig. 2).

**Fig. 2.**
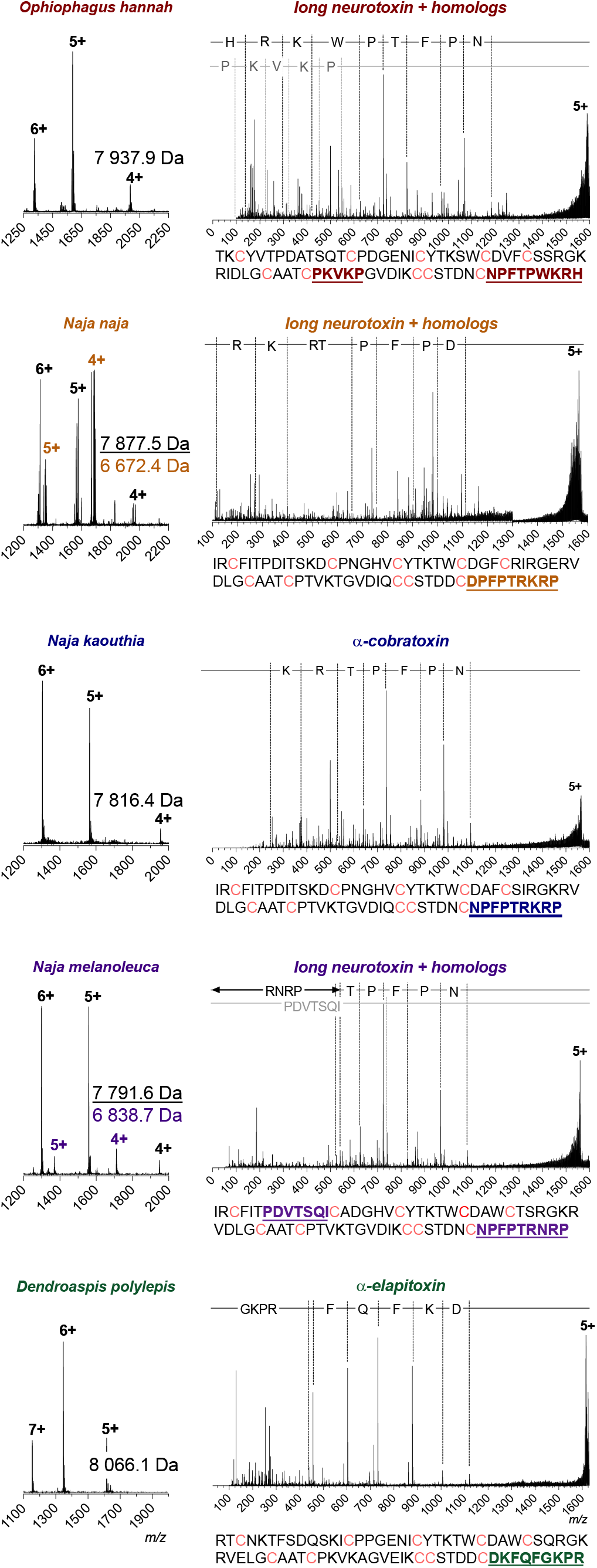
Intact masses and top-down sequence analysis of toxins bound by 2554_01_D11. Names above the mass spectra have been color-coded for each species as follows; *O. hannah* (red), *N. naja* (orange), *N. kaouthia* (blue), *N. melanoleuca* (purple) and *D. polylepis* (green). The spectra on the left-hand side show the charge state distribution for the toxins ejected from the antibody complex by applying a high cone voltage, where the masses of the identified toxins are given in Daltons. The top-down sequence spectra for the most prominent charge state of each toxin are shown in the right-hand side. The difference in *m/z* are outlined via dotted lines on top and match the specific amino acid or peptide. The full amino acid sequence for the proposed identity of the toxins is given below each spectrum, with the matching peptides found during the top-down analysis colored and underlined. Cysteines in the sequence are colored pink.

The sequences of the toxins bound by 2554_01_D11 were investigated via top-down proteomics to confirm the identities of these toxins. For these experiments, the toxin:antibody complexes were purified using SEC. Toxins were dissociated from the antibody by applying a high cone voltage. This is a focusing voltage applied to the cone, which is located in the source region of the instrument. Increasing this voltage leads to harsh condition that can dissociate noncovalent complexes. Since this dissociation occurs before the quadrupole, the most prominent charge state of each ejected toxin could then be isolated using MS/MS for top-down sequencing. This isolation is important, as it ensures that the peptide fragmentation peaks only correspond to the toxin of interest. For the toxins of masses between 7,800 and 8,200 Da, only one readily discernible peptide fragment series was detected for each precursor ion. The limited amount of sequence data obtained from these experiments is attributed to the presence of disulfide bonds present in snake venom toxins, which cannot be broken using this fragmentation technique^27–29^.

A BLAST search against all available elapid protein sequences revealed that the peptide sequences obtained by top-down analysis were unique to long chain α-neurotoxins and that each peptide only had one complete match to long chain α-neurotoxins homologs from the investigated venom. Sequence data combined with the detected masses of the toxins revealed that 2554_01_D11 was capable of binding to long chain α-neurotoxin-containing SEC fractions across all tested venoms. This suggested that this antibody is cross-reactive against long chain α-neurotoxins present in all five tested venoms, further highlighting the broadly cross-reactive behavior of 2554_01_D11.

The toxin homologs specifically identified to be bound by 2554_01_D11 were long neurotoxin 2 (A8N285) from *O. hannah*, α-cobratoxin (P01391) from *N. kaouthia*, long neurotoxin 2 (P01388) and long neurotoxin (P0DQQ2) from *N. melanoleuca*, long neurotoxin 4 (P25672) from *N. naja*, and α-elapitoxin (P01396) from *D. polylepis*. In addition, the antibody was shown to bind α-bungarotoxin (P60615) from *B. multicinctus* using SPR (data not shown). The average sequence similarity of the seven toxins was 62% (stdev: 9.9%), with an identity of 38% across all toxins; a total of 28 amino acid positions were identical across all toxins (Fig. 3A). The highest identity was observed between α-cobratoxin and long neurotoxin 2 from *N. melanoleuca* (83%) and the lowest identity was observed between long neurotoxin 2 from *N. melanoleuca* and α-bungarotoxin (51%). Additionally, a structural comparison was performed via root-mean-square deviation (RMSD) and revealed a mean pruned/total similarity of 0.81Å/3.1Å (stdev: 0.26Å/1.23Å), respectively; the best match appeared to be between long neurotoxin and long neurotoxin 2 from *N. melanoleuca* (0.23Å/0.23Å) and the worst match appeared to be between long neurotoxin 2 from *O. hannah* and α-cobratoxin (pruned: 1.18Å) and α-elapitoxin and α-bungarotoxin (total: 4.5Å; Fig 3B). For α-cobratoxin, the amino acid residues involved in binding to the nicotinic acetylcholine receptor have been highlighted both in the sequence (Fig. 3A) and in the structure on the toxin (Fig. 3C). Additionally, the residues that through a high-density peptide microarray-based study^30^ have been identified to be involved in the binding between antivenom-derived antibodies and α-elapitoxin and long neurotoxin 2 from *N. melanoleuca*, respectively, have been highlighted in the toxin sequence in Fig. 3A and in the toxin structure in Fig. 3D and 3E.

**Fig. 3.**
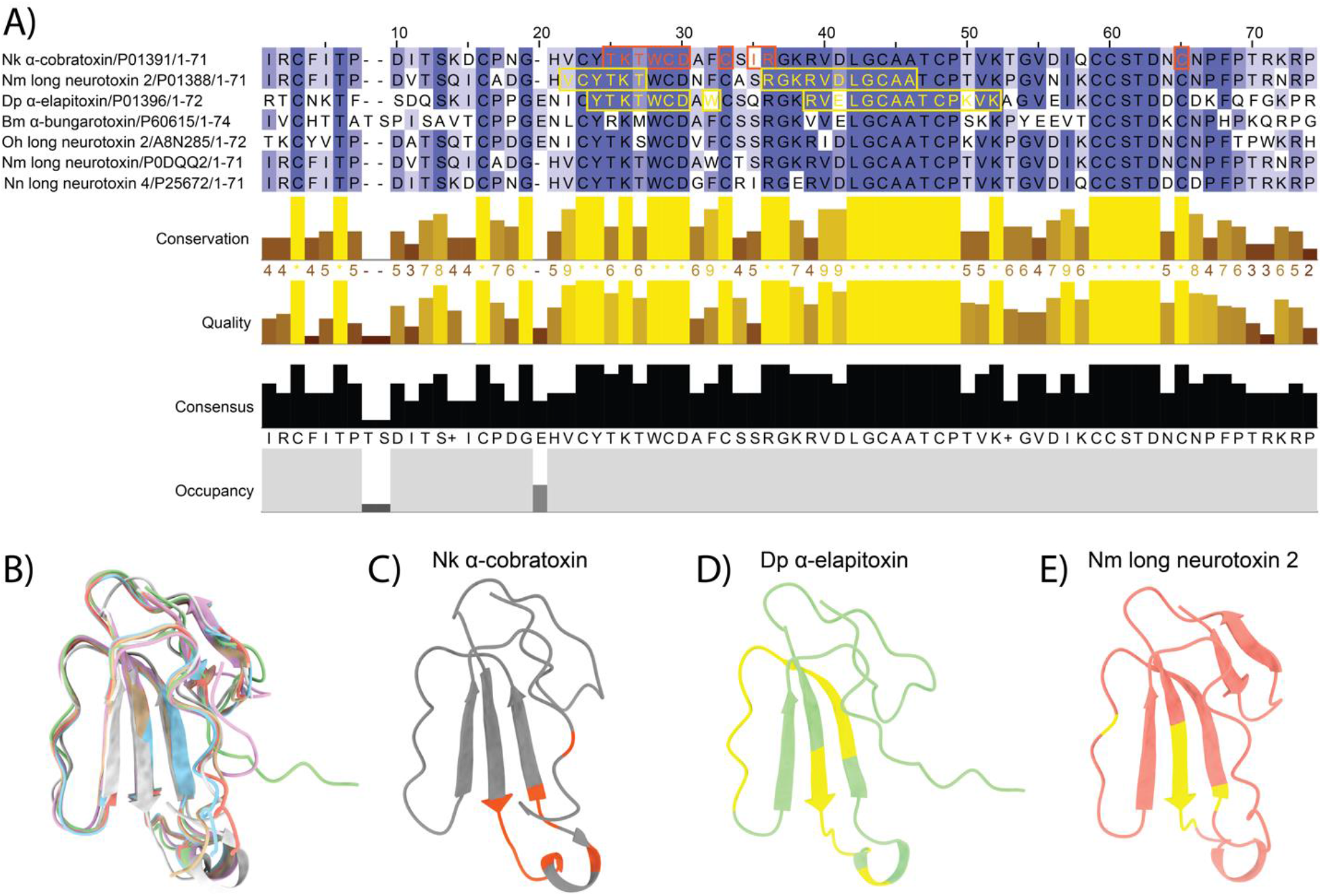
Alignment and epitope identification of all investigated long chain α-neurotoxins, *i.e*. α-cobratoxin (P01391/1CTX) from *N. kaouthia*, α-elapitoxin (P01396/AF-P01396) from *D. polylepis*, α-bungarotoxin (P60615/1HC9) from *B. multicinctus*, long neurotoxin 2 (A8N285/AF-A8N285) from *O. hannah*, and long neurotoxin (P0DQQ2), long neurotoxin 4 from *N. naja* (P25672/AF-P25672), and long neurotoxin 2 (P01388/AF-P01388) from *N. melanoleuca*. A) Sequence alignment using Clustal Omega with boxes indicating residues involved in binding to the nicotinic acetylcholine receptor (orange) or bound by antivenom antibodies (yellow). B) Structural alignment in ChimeraX with the following colors representing each toxin: orange (long neurotoxin 2 from *N. melanoleuca*), beige (long neurotoxin 2 from *O. hannah*), purple (α-bungarotoxin), green (α-elapitoxin from *D. polylepis*), blue (long neurotoxin 4 from *N. naja*), and grey (α-cobratoxin from *N. kaouthia*). C) Amino acid residues on α-cobratoxin known to be involved in binding to its native target, *i.e*., the nicotinic acetylcholine receptor^31^ (orange). D) Amino acid residues in α-elapitoxin suggested to be bound by antivenom antibodies based on high-density peptide microarray analysis^30^ (yellow). E) Amino acid residues in long neurotoxin 2 from *N. melanoleuca* suggested to be bound by antivenom antibodies based on high-density peptide microarray analysis^30^ (yellow).

### Increased *in vitro* neutralization potency and broadening of cross-neutralization

After having established the broadly cross-reactive nature of one of the top two chain-shuffled antibodies (2554_01_D11), automated patch-clamp technology was applied to assess if binding translated into functional neutralization *in vitro* for 2551_01_A12 and 2554_01_D11, as well as for the parental clone, 368_01_C05. Here, a human derived cell line endogenously expressing the nAChR was used for measuring the acetylcholine-dependent current. α-cobratoxin inhibited this current in a concentration-dependent manner, and the IC80 value for the toxin was determined. Thereafter, the concentration-dependent neutralization of the current-inhibiting effect of α-cobratoxin by the three antibodies was determined. Results demonstrated that all three antibodies were able to fully neutralize the effects of α-cobratoxin, whereas an irrelevant isotype control antibody (recognizing a dendrotoxin) had no effect (Fig. 4A). The parental antibody, 368_01_C05 neutralized α-cobratoxin mediated inhibition of acetylcholine-dependent currents with an EC50 value of 4.9 nM and relatively shallow concentration-response curve slope. In contrast, the optimized antibodies, 2551_01_A12 and 2554_01_D11 exhibited improved EC50 values of 2.6 nM and 1.7 nM respectively with steeper slopes for the concentration-response curves. These EC50 values translate into toxin:antibody molar ratios of 1:1.23 for 368_01_C05, 1:0.65 for 2551_01_A12, and 1:0.43 for 2554_01_D11. Since each IgG has two binding sites, the theoretically lowest amount of IgG needed to neutralize the effect of one toxin would be 0.5 IgGs.

**Fig. 4.**
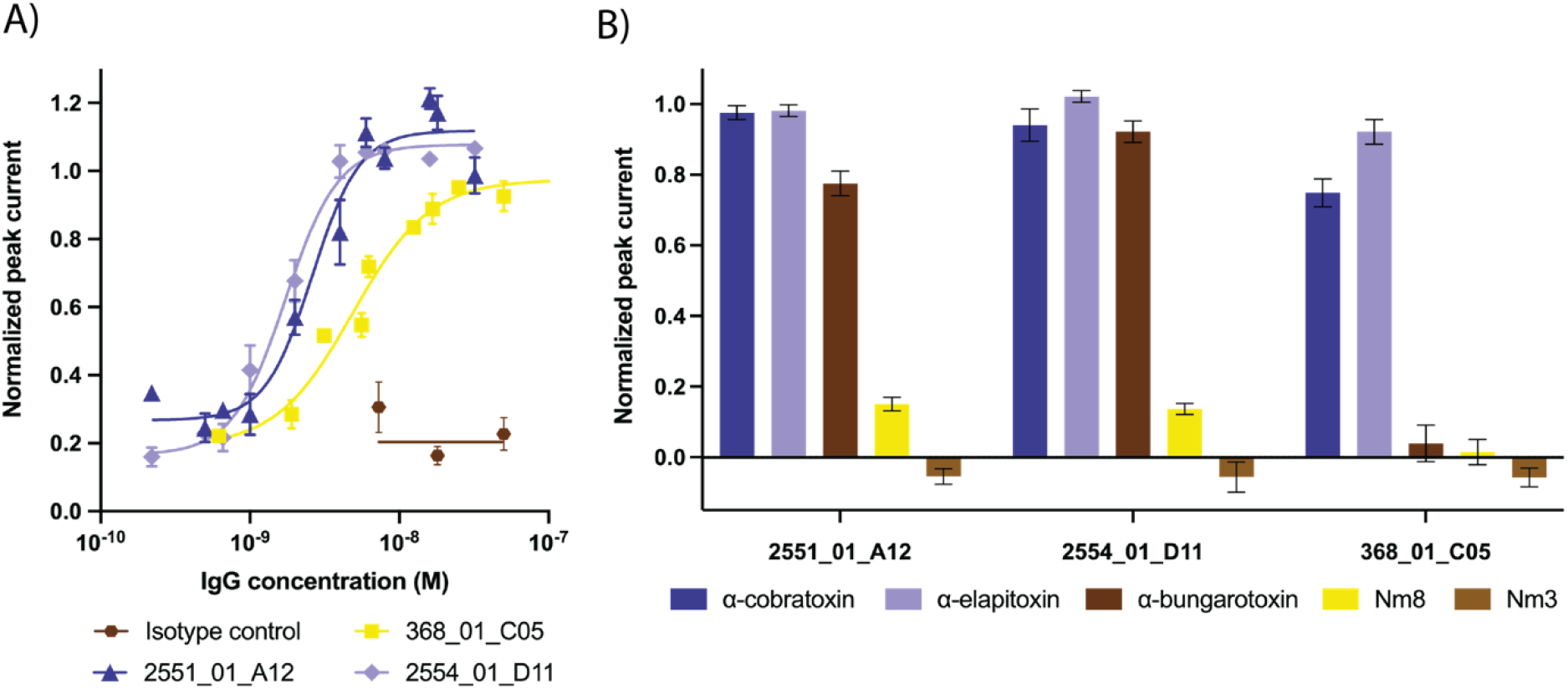
Electrophysiological determination of the *in vitro* cross-neutralizing potential of 2551_01_A12, 2554_01_D11, and 368_01_C05. Automated patch-clamp experiments were performed to determine the ability of the antibodies to prevent the current-inhibiting effect that α-neurotoxins exert on the nAChR. A) Concentration-response curves illustrating how increasing concentrations of the three antibodies prevent nAChR blocking by α-cobratoxin. B) Single concentration plot outlining the cross-neutralizing potential of the antibodies against α-cobratoxin from *N. kaouthia*, α-elapitoxin from *D. polylepis*, α-bungarotoxin from *B. multicinctus*, and Nm8 from *N. melanoleuca*. In addition, a negative control Nm3, a fraction from *N. melanoleuca* venom containing a short α-neurotoxin, was included. The toxin to antibody molar ratios used were 1:22 for α-cobratoxin, 1:40 for α-elapitoxin, 1:5 for α-bungarotoxin, 1:2.3 for Nm8, and 1:3.2 for Nm3.

To determine if the increased cross-reactivity to other α-neurotoxins translated into cross-neutralization, a single concentration antibody screen was set up using the Qube 384 system. Here, the three antibodies (368_01_C05, 2551_01_A12, and 2554_01_D11) were tested against α-cobratoxin from *N. kaouthia*, α-elapitoxin from *D. polylepis*, and Nm8 from *N. melanoleuca*, which all were toxins that 2554_01_D11 had been shown to bind through native MS. In addition, α-bungarotoxin was included, as it has 58% sequence identity to α-cobratoxin, is commercially available, and is an important toxin to neutralize in the venom of *B. multicinctus*. As a control, Nm3, a venom fraction from *N. melanoleuca* containing a short chain α-neurotoxin that also binds to the nAChR, but is not bound by any of the three antibodies, was included. This automated patch-clamp screening revealed that α-cobratoxin and α-elapitoxin could be neutralized by all three antibodies in this assay (Fig. 4B). Additionally, the chain-shuffled clones were able to neutralize α-bungarotoxin and partially neutralize the α-neurotoxins present Nm8, none of which was achieved by the parental clone. Collectively, the results of the *in vitro* neutralization assays using automated patch-clamp demonstrated that the chain-shuffled antibodies were both more potent in their neutralization of α-cobratoxin, as well as more broadly neutralizing than the parental antibody, inhibiting the effect of α-neurotoxins from snakes of three different genera inhabiting both Asia and Africa. Based on binding, developability, affinity, expression, and *in vitro* neutralization data, 2554_01_D11 was selected as the top candidate for *in vivo* testing.

### *In vitro* neutralization data translates to complete or partial *in vivo* neutralization of snake venoms from different genera and continents

To verify that the *in vitro* cross-neutralization potential of 2554_01_D11 translated into *in vivo* cross-neutralization, animal experiments were set up to evaluate the ability of the antibody to prevent or delay venom-induced lethality. Here, snake venoms from three different species belonging to three genera, one from Africa, *i.e. D. polylepis*, and two from Asia, *i.e. N. kaouthia* and *O. hannah*, were included. Notably, each of these venoms contain a substantial amount of long chain α-neurotoxins (a relative abundance of 13.2%^18^, 55%^17^, and ~20%^32^, respectively). Two LD_50_s of each venom were preincubated with 2554_01_D11 in a 1:1 and 1:2 toxin:antibody molar ratio for *N. kaouthia* and *O. hannah* or a 1:3 toxin:antibody molar ratio for *D. polylepis* before being administered i.v. to the mice. As controls, mice were injected with venom alone, venom preincubated with commercial antivenoms (except in the case of *O. hannah*, where no antivenom was available), or venom preincubated with an antibody isotype control.

Results of the studies demonstrated that all mice in the venom only control group, as well as the mice receiving venom preincubated with the isotype control antibody, died within the first hour after the challenge, with evident signs of limb paralysis and respiratory difficulty. As expected, mice receiving *N. kaouthia* or *D. polylepis* venoms preincubated with commercially available antivenoms survived for the entire observation period, and no signs of neurotoxicity were observed. In experiments where mice were injected with venoms incubated with 2554_01_D11, results varied depending on the venom. In the case of *N. kaouthia* venom, complete neutralization was observed at both toxin:antibody molar ratios, since mice survived during the 48-hour observation time. Moreover, mice did not show any signs of neurotoxicity, *i.e*., limb paralysis or respiratory difficulty along the whole period. In the case of *O. hannah*, there was a dose-dependent delay in the time of death, as compared to controls receiving venom alone. Likewise, a delay in the time of death was observed in the case of *D. polylepis* venom (Fig 5).

**Fig. 5.**
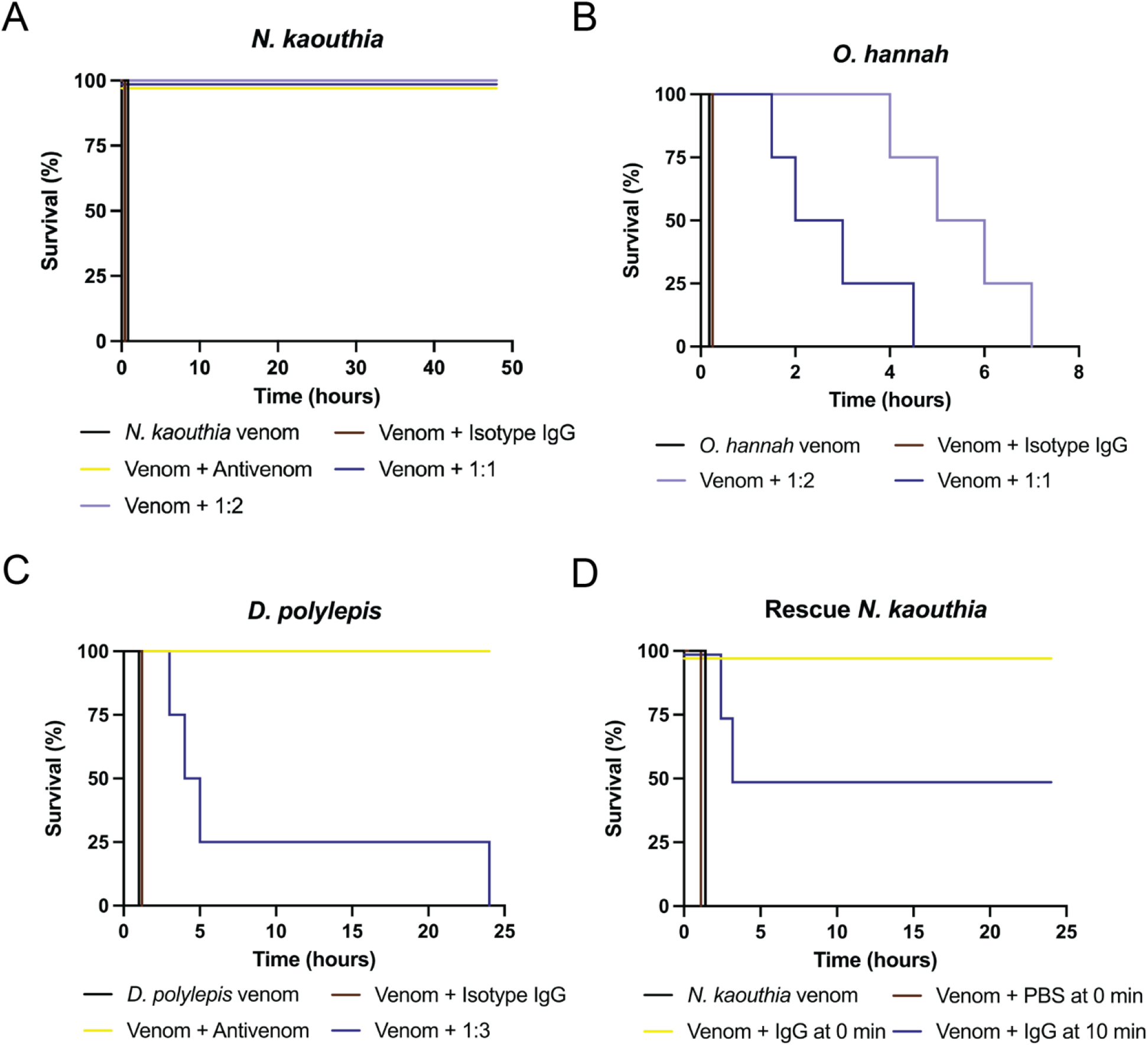
Kaplan-Meier survival curves for the antibody 2554_01_D11. A), B), and C) Mixtures containing 2 LD_50_s of venom of either *N. kaouthia*, *O. hannah*, or *D. polylepis* were preincubated with the antibody, 2554_01_D11, at various toxin:antibody ratios and then administered i.v. to groups of four mice. Controls included mice receiving venom alone or venom incubated with either an irrelevant isotype antibody control or a commercial horse-derived antivenom. Signs of toxicity were observed, and deaths were recorded for a maximum period of 48 hours. D) 2 LD_50_s *N. kaouthia* venom was administered s.c. following i.v. administration of IgG 2554_01_D11 either immediately following venom administration or 10 minutes after venom administration. As a control, mice were administered venom s.c. and PBS i.v. Signs of toxicity were observed, and deaths were recorded for 24 hours.Mice receiving antibody immediately following venom administration had a dose of 1:2.5 toxin to antibody molar ratio while mice receiving antibody 10 minutes post venom administration had an antibody dose of 1:2 toxin to antibody molar ratio.

Next, the ability of the antibody to abrogate lethality of *N. kaouthia* venom in rescue-type experiments was assessed. For this, the subcutaneous (s.c.) route of venom injection was used in order to better reproduce the actual circumstances of envenoming. The estimated LD_50_ by the s.c. route was 10.3 μg (95% confidence interval: 5.0 – 16.8 μg). Mice were challenged by the s.c. route with a dose of venom corresponding to 2 LD_50_s, *i.e*., 20 μg, followed by the i.v. administration of the 2554_01_D11 antibody in a volume of 100 μL, at a molar ratio of 1:2.5 (toxin:antibody). Control mice injected with venom only died within 40 – 60 min, with evident signs of limb and respiratory paralysis. When the antibody was administered immediately after venom injection, all mice survived the 24-hour observation period and did not show any evidence of limb or respiratory paralysis. When the antibody was provided 10 min after venom injection, two out of four mice died, but there was a delay in the time of death (150 min and 180 min). The other two mice survived the 24-hour observation time and did not show signs of paralysis (Fig 5D).

## DISCUSSION

Here, we demonstrate that an antibody discovered from a naïve human library with limited cross-reactivity to other α-neurotoxins and without the ability to prevent lethality induced by α-cobratoxin in mice, can be improved by light chain-shuffling, resulting in enhanced affinity, potency, and cross-neutralization capacity.

The most promising antibody we discovered, 2554_01_D11, bound seven long chain α-neurotoxins arising from five snakes of four genera distributed across both Asia and Africa. Notably, cross-reactive binding was detected despite sequence alignment of the seven α-neurotoxins revealing substantial differences in sequence identity, with an overall identity of only 31%. This is likely due to the fact that, despite the low overall identity, a total of 29 positions in the toxin sequences contain identical amino acids across all seven α-neurotoxins, including most of the residues previously identified as playing a significant role in the binding between α-cobratoxin/α-bungarotoxin and the nAChR^31^. Specifically, these amino acid residues include Trp25, Cys26, Asp27, Ala28, Phe29, Cys30, Arg33, Lys35, and Arg36/Val39 (α-cobratoxin/α-bungarotoxin) on loop II and Phe65/Val39 (α-cobratoxin/α-bungarotoxin) on the C-terminus, where a single mutation of one of these residues has been shown to cause a more than five-fold decrease in affinity to the nAChR^33^. Furthermore, high-density peptide microarray analysis previously suggested that positions 22-27 and 36-46 represent linear B-cell epitopes for antibodies to long neurotoxin 2 from *N. melanoleuca*, recognized by one of the most effective antivenoms available (SAIMR, produced by SAVP), like positions 24-30, 32, and 39-50 do for α-elapitoxin^30^. This emphasizes the potential importance of Trp25, Cys26, Asp27, Ala28, Phe29, Cys30, and Arg36 for the ability of antibodies to recognize this toxin. Together, this presents a plausible explanation for the broad cross-reactivity we observed for 2554_01_D11 and indicates the importance of epitope similarity (as opposed to overall sequence identity) in the pursuit of cross-reactive antibodies^4^. However, the boundaries of the cross-reactivity of 2554_01_D11 have not been established in this study, as all long chain α-neurotoxins investigated were recognized by the antibody. Future work aiming to investigate the boundaries of cross-reactivity could include testing the antibody for binding to long chain α-neurotoxins from other snakes, such as *B. candidus*^34^, *B. fasciatus*^34^, or *N. haje legionis*^35^. Such studies could potentially provide general cues to how antibody cross-reactivity can be optimized for antibodies targeting toxins and similar antigens.

Besides cross-reactive binding, we also demonstrated the broad neutralizing potential of 2554_01_D11. *In vivo* studies showed that lethality induced by three snake venoms of different genera distributed across both Asia and Africa was either prevented or delayed by the antibody, 2554_01_D11. For *N. kaouthia* venom, the antibody was able to completely prevent lethality in envenomed mice with no signs of neurotoxicity even at the lowest tested toxin:antibody molar ratio of 1:1. Moreover, this antibody was able to neutralize lethality induced by the venom of *N. kaouthia* in rescue-type experiments, which more closely resemble the actual circumstances of envenoming^36^. Complete neutralization was achieved when the antibody was administered immediately after the venom challenge, and even after a delay of 10 min between venom and antibody administration, 2 out of 4 mice survived and death was delayed in the other two.

Despite the antibody 2554_01_D11 possessing very similar affinities to α-cobratoxin and α-elapitoxin from *D. polylepis*, the antibody was unable to prevent lethality induced by *D. polylepis* venom, even at the tested toxin:antibody molar ratio of 1:3. Survival was, however, prolonged several hours, suggesting that the antibody did provide partial neutralization of the venom. These results are perhaps not surprising, as the venom of *D. polylepis* is more complex than that of *N. kaouthia*, and it is well-established that toxins other than long chain α-neurotoxins (*i.e*. short chain α-neurotoxins and dendrotoxins) play important roles for the toxicity of *D. polylepis* venom^18^. Where lethality of *N. kaouthia* venom is mainly attributed to the high content of long chain α-neurotoxins, the short chain α-neurotoxins present in *D. polylepis* venom have been estimated to contribute with about a third of the toxicity of the venom^18^. Thus, even if all long chain α-neurotoxins were neutralized in the venom, neutralization of short chain α-neurotoxins and possibly even dendrotoxins could still be necessary to prevent venom-induced lethality. As the affinity of the antibody to α-elapitoxin was almost identical to that of α-cobratoxin, we speculate that the antibody, 2554_01_D11, might be able to neutralize the effects of the long chain α-neurotoxins from *D. polylepis*, but that the mice eventually die due to lethality induced by short chain α-neurotoxins and possibly dendrotoxins. Similarly, the lethal effects of the venom of *O. hannah* were only delayed, probably since this venom also consists of a mixture of both long and short chain α-neurotoxins.

In this study, monoclonal IgG antibodies with broad cross-reactivity to different long chain α-neurotoxins were discovered, and an epitope binning study revealed that the cross-reactive antibody 2554_01_D11 bound the same or an overlapping epitope to the previously reported antibody 2552_02_B02, which only recognizes α-cobratoxin^8^. In the future, determination of the structure of the two antibodies in complex with α-cobratoxin might provide further insight into how two antibodies binding to the same or overlapping epitope with similar affinity can display such different levels of cross-reactivity.

When comparing the neutralizing capacities of the two antibodies, 2554_01_D11 and 2552_02_B02, another noteworthy observation emerges. Antibody 2554_01_D11 possessed significantly higher efficacy in neutralizing *N. kaouthia* venom *in vivo* when compared to what was reported for 2552_02_B02^8^. Where 2554_01_D11 neutralized all signs of neurotoxicity at the lowest tested dose of 1:1 toxin to antibody molar ratio, 2552_02_B02 only prevented lethality induced by *N. kaouthia* venom in 3 out of 4 mice at a 1:4 molar ratio, with mice showing clear signs of neurotoxicity. These results were especially remarkable, as the two antibodies performed similarly both in electrophysiological *in vitro* neutralization assays and had similar affinities to α-cobratoxin (490 pM for 2552_02_B02 and 1.78 nM for 2554_01_D11). Except for the difference in cross-reactivity profiles, the only significant difference between the two antibodies was in their developability profiles. In these developability assessment assays, 2554_01_D11 performed similarly to Aliricumab (control for good developability), whereas 2552_02_B02 performed comparably to Bococizumab (control for poor developability)^25^. It is thus possible that this difference in self-association and interaction with the SEC column seen in these assays may correlate with different pharmacokinetic or pharmacodynamic properties of the two antibodies, which could explain their contrasting performance *in vivo*. This thus suggests that detailed developability characterization should be included as part of early discovery to maximize the *in vivo* efficacy and clinical success of recombinant antivenoms.

In conclusion, this study demonstrates the utility of combining cross-panning strategies in phage display with affinity maturation using chain-shuffling for the development of high-affinity human monoclonal IgG antibodies that show broadly-neutralizing effects against neurotoxic elapid snake venoms *in vitro* and *in vivo*. Such antibodies might be useful for designing future envenoming therapy, but more importantly, the pipeline presented here could also be exploited for the development of broadly-neutralizing antibodies against other targets of medical importance. These targets could include toxins from other venomous animals than snakes, but also hypervariable and mutating antigens from infectious bacteria, viruses, and parasites, or even neoepitopes in non-infectious diseases.

## MATERIALS AND METHODS

### Toxin preparation

α-cobratoxin (L8114), α-bungarotoxin (L8115), and whole venoms from *N. kaouthia* (L1323), *N. melanoleuca* (L1318), *D. polylepis* (L1309), and *O. hannah* (L1357) were obtained from Latoxan SAS, France. Venom fractions containing long α-neurotoxins (Dp7 from *D. polylepis* and Nm8 from *N. melanoleuca*) were isolated from crude venom by fractionation using RP-HPLC (Agilent 1200). Venoms were fractionated using a C18 column (250 × 4.6 mm, 5 μm particle; Teknokroma) and elution was carried out at 1 mL/min using Solution A (water supplemented with 0.1% TFA) and a gradient towards solution B (acetonitrile supplemented with 0.1% TFA): 0% B for 5 min, 0–15% B over 10 min, 15–45% B over 60 min, 45–70% B over 10 min, and 70% B over 9 min^18,24^. Fractions were collected manually and evaporated using a vacuum centrifuge. Toxins were dissolved in phosphate buffered saline (PBS) and biotinylated using a 1:1 (toxin:biotinylation reagent) molar ratio for α-cobratoxin and 1:1.5 molar ratio for the remaining toxins as described elsewhere^6^. Following biotinylation, toxins were purified as well as the degree of biotinylation was determined as previously described^8^.

### Library generation using chain-shuffling

Light chain-shuffled libraries containing the heavy chain of antibody 368_01_C05 were created as described previously^8^. Two libraries were created, one with kappa and one with lambda light chains. The size of the kappa library was 1.67 × 10^8^ whereas the lambda library was 1.01 × 10^8^. Colony PCR revealed 96% of the transformants to have successful heavy chain ligation.

### Library rescue, solution-based phage display selection, and polyclonal DELFIA

Phages were rescued from the light chain-shuffled libraries and three rounds of selections were performed as described elsewhere^37^ with a few exceptions. Phages were not concentrated using PEG precipitation, but instead phage-containing supernatants were blocked in PBS supplemented with Milk Protein (MPBS) and used directly for selections. Additionally, deselection of streptavidin-specific phages was performed as described previously^8^. Lastly, to obtain scFvs with high affinity and broad cross-reactivity to different long α-neurotoxins cross-panning between α-cobratoxin and Dp7 was performed as previously described^7^ as well as antigen concentrations were lowered between each round of selection starting at 10 nM and ending at 4 pM. The kappa and lambda libraries were mixed before the first round of selections. To assess the polyclonal output of the selections a polyclonal DELFIA was performed determining binding to both α-cobratoxin, Dp7, and MPBS.

### Subcloning, screening, and sequencing of scFvs

The genes encoding the scFvs from five of the obtained selection outputs (representing different cross-panning strategies) were subcloned into the pSANG10-3F expression vector to allow for monoclonal screening of the clones as described elsewhere^6^. Briefly, scFv-encoding genes from five selected output phage libraries were amplified using primers M13leadseq (AAATTATTATTCGCAATTCCTTTGGTTGTTCCT) and Notmycseq (GGCCCCATTCAGATCCTCTTCTGAGATGAG) before restriction using *Nco*I and *Not*I restriction endonuclease sites. The genes were ligated into pSANG10-3F and transformed into *E. coli* strain BL21 (DE3) (New England Biolabs). From each selection output 184 clones were selected and expressed in 96-well plates. The scFvs were evaluated for their binding to α-cobratoxin, Dp7, streptavidin, and milk protein using a monoclonal DELFIA assay as described previously^8^. From this, 329 clones were cherry-picked and sequenced (Eurofins Genomics sequencing service) using S10b primer (GGCTTTGTTAGCAGCCGGATCTCA). The antibody frameworks and the CDR regions of the light chains were annotated using Geneious Biologics (Biomatters), and 67 clones were identified as unique based on light chain CDR3 regions.

### Reformatting to IgG and Fab format

A total of 62 clones were selected for reformatting into the fully human IgG1 and Fab format. The reformatting into the IgG1 format was completed as described in Laustsen *et al*.^6^, whereas reformatting into the Fab format was completed as described in Ledsgaard *et al*^8^. The binding of the IgGs to α-cobratoxin, Dp7, Nm8, and streptavidin was assessed and ranked using an expression-normalized capture (ENC) assay described previously^8^.

### Developability characterization

To aid in the selection of the top antibody candidates for further characterization, the biophysical behavior of the 62 reformatted clones in the IgG format was characterized using HPLC-SEC and AC-SINS. For HPLC-SEC, the purified antibodies were loaded onto a Superdex 200 Increase 5/150 column at a flow rate of 0.25 mL/min using an Agilent 1100 HPLC instrument. AC-SINS was performed as described in Liu *et al*.^38^ with the modifications described in Dyson *et al*.^25^.

### Selection, expression, and purification of IgGs

Based on ranking and cross-reactivity in IgG binding in ENC assay, IgG expression yield, HPLC-SEC profile, AS-SINS shift, and sequence diversity six antibodies (2551_01_A12, 2551_01_B11, 2554_01_D11, 2555_01_A01, 2555_01_A04, and 2558_02_G09) were selected for further characterization. The IgGs were expressed and purified as previously described^6^.

### Surface plasmon resonance

The binding affinity of the corresponding Fab versions of the top six affinity matured antibodies as well as the parental clone to α-cobratoxin and Dp7 was determined using Surface Plasmon Resonance (SPR) (BIAcore T100, GE Healthcare). Antigen immobilization and affinity measurements were performed as previously described^8^. Based on affinity measurements the top two clones for further characterization were selected to be 2551_01_A12 and 2554_01_D11.

Epitope binning experiments were performed using a sandwich setup, whereby one antibody was immobilized on a CM5 sensor chip using amine coupling, prior to flowing α-cobratoxin with a competing antibody. The 2554_01_D11 Fab (10 μg/mL, 10mM sodium acetate pH5.0) was immobilized to a level of 450RU, and 20nM α-cobratoxin (10mM, HEPES, 150 mM NaCl, 50 mM MES, 0.05% P20, pH7.4) was incubated with 200nM of either test 2552_02_B02 Fab or control Fab. Dual binding was then measured by injecting the α-cobratoxin and Fab solution over the immobilized 2554_01_D11 flow cell for 120s. A blank flow cell was used as a reference, and was subtracted from the test flow cell for analysis.

### Determining cross-reactivity using native mass spectrometry

#### Sample preparation

Venoms and antibody samples were fractionated and exchanged into 200 mM ammonium acetate by size exclusion chromatography (SEC) as previously described^39,40^. These experiments were performed on a Superdex Increase 200 10/300 GL column (Cytiva, Massachusetts, United States) pre-equilibrated with 200 mM ammonium acetate. Samples were collected and stored a 4 °C until used. Prior to analysis, aliquots of the venom and IgG 2554_01_D11 SEC fractions were mixed in a 1:1 ratio (*v*/*v*). The final concentration of the antibody was approximately 3 μM after mixing. The concentration of toxins in the SEC fractions were not adjusted prior to mixing with the antibody.

#### Native mass spectrometry

All mass spectrometry (MS) experiments were performed on a SELECT SERIES cyclic IMS mass spectrometer (Waters, Manchester, U.K.) which was fitted with a 32,000 *m/z* quadrupole, as well as an electron capture dissociation (ECD) cell (MSvision, Almere, Netherlands), the latter of which was situated in the transfer region of this mass spectrometer. Approximately 4 μL of sample were nano-sprayed from borosilicate capillaries (prepared in-house) fitted with a platinum wire. Spectra were acquired in a positive mode, with the *m/z* range set to 50-8,000. Acquisitions were performed for five minutes at a rate of 1 scan per second. The operating parameters for the MS experiments were as follows, unless otherwise stated: capillary voltage, 1.2 - 1.5 kV; sampling cone, 20 V; source offset, 30 V; source temperature, 28 °C; trap collision energy, 5 V; transfer collision energy, 5 V; Ion guide RF, 700 V. This instrument was calibrated with a 50:50 acetonitrile:water solution containing 20 μM cesium iodide (99.999%, analytical standard for HR-MS, Fluka, Buchs, Switzerland) each day prior to measurements.

#### Top-down proteomics of toxins bound by 2554_01_D11

The toxin:antibody complexes were purified using SEC, using the methods described above. Toxins were ejected from the protein complex during the MS experiments by setting the cone voltage to 160 V. The 5^+^ ions (most abundant charge state) of the ejected toxins were selected by tandem MS (MS/MS) and subjected to fragmentation by applying a trap voltage between 80 and 100 V as well as a transfer voltage between 20 and 50 V. Peptide sequence assignment was performed for 1^+^ fragmentation ions using the BioLynx package, which is a part of the MassLynx v4.1 software.

### Sequence alignment

Sequence alignment was performed in Clustal Omega^41^ and visualised in Jalview^42^ using α-cobratoxin (P01391) from *N. kaouthia*, α-elapitoxin (P01396) from *D. polylepis*, α-bungarotoxin (P60615) from *B. multicinctus*, long neurotoxin 2 (A8N285) from *O. hannah*, and long neurotoxin (P0DQQ2) and long neurotoxin 2 (P01388) from *N. melanoleuca*. Structures for each toxin were retrieved prioritising high-resolution X-ray resolved structures and included the following: P01391 = 1CTX (2.8Å, X-ray), P01388 = AF-P01388-F1 (AlphaFold2 predicted), P01396 = AF-P01396-F1 (AlphaFold2 predicted), P60615 = 1HC9 (1.8Å, X-ray), A8N285 = AF-A8N285-F1 (AlphaFold2 predicted), and P0DQQ2 = AF-P0DQQ2-F1 (AlphaFold2 predicted). Structural alignment and root-mean-square deviation (RMSD) analysis were performed in ChimeraX^43^. Epitopes of P01388 and P01396 were identified using the STAB Profiles tool^30^ (https://venom.shinyapps.io/stab_profiles/).

### *In vitro* neutralization using electrophysiology (QPatch)

To determine the ability and potency with which the affinity matured clones 2551_01_A12 and 2554_01_D11, as well as the parental antibody 368_01_C05, were able to neutralize the effects of α-cobratoxin, whole-cell patch-clamp experiments were conducted using Rhabdomyosarcoma cells (ACTT) as previously described^8^. In brief, the nAChR-mediated current elicited by acetylcholine was determined in the presence of 4 nM α-cobratoxin or 4 nM α-cobratoxin preincubated with different concentrations of the three IgGs on a QPatch II automated electrophysiology platform (Sophion Bioscience). As a control a dendrotoxin-specific IgG was included. The inhibitory effect of α-cobratoxin was normalized to the full acetylcholine response and a non-cumulative concentration-response plot was plotted. A Hill fit was used to obtain IC50 values for each of the three IgGs.

### *In vitro* cross-neutralization using electrophysiology (Qube 384)

To determine the broader cross-neutralizing potential of the top two affinity matured antibodies and the parent antibody, automated patch-clamp experiments using the Qube 384 electrophysiology platform (Sophion Bioscience) were conducted. The three IgGs (32.5 nM) were tested against α-cobratoxin (1.47 nM), α-bungarotoxin (6.5 nM), Dp7 (0.81 nM), Nm8 (14 nM), Nm3 (10.3 nM). A dendrotoxin-specific IgG was included as a control.

### Animals

Animal experiments were conducted in CD-1 mice of both sexes weighing 18-20 g. Mice were supplied by Instituto Clodomiro Picado and experiments were conducted following protocols approved by the Institutional Committee for the Use and Care of Animals (CICUA), University of Costa Rica (approval number CICUA 82-08). Mice were provided food and water *ad libitum* and housed in plastic cages in groups of 4.

### *In vivo* preincubation experiments

The *in vivo* cross-neutralizing potential of 2554_01_D11 against whole venoms of *N. kaouthia, D. polylepis*, and *O. hannah* was assessed by i.v. injection of IgG preincubated with venom using groups of four mice per treatment. Mixtures of a constant amount of venom and various amounts of antibody were prepared and incubated for 30 min at 37 °C. Then, aliquots of the mixtures, containing 2 LD_50_s of venoms (for *N. kaouthia* 9.12 μg, for *D. polylepis* 25.8 μg, and for *O. hannah* 40 μg) were injected in the caudal vein of mice using an injection volume of 150-200 μL. Control mice were injected with either venom alone, venom preincubated with an isotype control IgG or, in the cases of *N. kaouthia* and *D. polylepis* venoms, they were preincubated with commercial horse-derived antivenoms. For *N. kaouthia* Snake Venom Antiserum from VINS Bioproducts Limited (Batch number: 01AS13100) was used at a ratio of 0.2 mg venom per mL antivenom. For *D. polylepis* Premium Serum and Vaccines antivenom (Batch number: 062003) was used at a ratio of 0.12 mg venom per mL antivenom.

The IgG was injected using 1:1 and 1:2 α-neurotoxin:IgG molar ratio for *N. kaouthia* and *O. hannah* and 1:3 α-neurotoxin:IgG molar ratio for *D. polylepis*. For calculating molar ratios, based on toxicovenomic studies, it was estimated that 55% *of N. kaouthia* venom^17^, 13.2% of *D.polylepis* venom^18^, and 40% of *O. hannah* consisted of α-neurotoxins^44^. Following injection, animals were observed for signs of neurotoxicity, and survival was monitored for 48 hours. Results were presented in Kaplan-Meier plots.

### *In vivo* rescue-type experiments

In order to assess whether the antibody was capable of neutralizing the venom of *N. kaouthia* in an experimental setting that more closely resembles the actual circumstances of envenoming, a rescue-type experiment was designed. For this, the s.c. route was used for injection of venom, while the antibody was administered i.v. First, the s.c. LD_50_ of *N. kaouthia* venom was estimated by injecting various doses of venom diluted in 100 μL of PBS into groups of four mice. Animals were observed during 24 hr, deaths were recorded and the LD_50_ was estimated by probits^45^. For neutralization experiments, groups of four mice first received a challenge dose of a challenge dose of 20.6 mg of venom by the s.c. route, corresponding to 2 LD_50_s. Then, mice receiving antibody 2554_01_D11 immediately following venom administration, 535 μg of antibody was administered i.v. in the caudal vein (in a volume of 100 μL) corresponding to a 1:2.5 toxin to antibody molar ratio. For mice receiving antibody 10 min after envenoming, 412 μg of the 2554_01_D11 antibody were administered i.v. in the caudal vein (in a volume of 100 μL), corresponding to a toxin to antibody molar ratio of 1:2. Mice were observed for the onset of neurotoxic manifestations and times of death were recorded and presented in Kaplan-Meier plots.

## Supplementary material

**Table S1.** Expression and developability data for 61 IgGs and parental IgG. The expression yield is provided for cultures in 96-well format. AC-SINS shift reflects IgG self-association propensity and SEC analysis provides information on elution volume as well as percentage of IgG monomer and dimers detected.

| Antibody ID | Production | AC-SINS | SEC analysis |  |  |
| --- | --- | --- | --- | --- | --- |
|  | Yield (mg/L) | Shift (nm) | Elution (mL) | Monomer (%) | Dimer (%) |
| 2551_01_A12 | 7,7 | 3 | 1,48 | 100,0 | 0,0 |
| 2554_01_D11 | 14,0 | 1 | 1,48 | 94,8 | 5,2 |
| 2552_02_B07 | 10,7 | 1 | 1,47 | 97,1 | 2,9 |
| 2558_02_G09 | 21,7 | 1 | 1,48 | 95,6 | 4,4 |
| 2551_01_A11 | 10,6 | 2 | 1,50 | 95,2 | 4,8 |
| 2554_01_E01 | 15,5 | 2 | 1,49 | 96,6 | 3,4 |
| 2551_01_B11 | 11,1 | 1 | 1,50 | 96,2 | 3,8 |
| 2554_02_F10 | 23,6 | 1 | 1,48 | 96,5 | 3,5 |
| 2551_01_A02 | 11,3 | 1 | 1,46 | 97,0 | 3,0 |
| 2552_01_G02 | 11,7 | 2 | 1,49 | 96,6 | 3,4 |
| 2554_01_E10 | 12,2 | 1 | 1,47 | 95,2 | 4,8 |
| 2554_02_F11 | 13,9 | 1 | 1,48 | 94,4 | 5,6 |
| 2555_01_A04 | 8,7 | 1 | 1,47 | 100,0 | 0,0 |
| 2555_01_A01 | 16,0 | 1 | 1,47 | 96,7 | 3,3 |
| 2554_02_G09 | 14,8 | 4 | 1,48 | 97,0 | 3,0 |
| 2558_01_E06 | 15,2 | 1 | 1,47 | 96,3 | 3,7 |
| 2554_01_E03 | 14,5 | 1 | 1,48 | 96,6 | 3,4 |
| 2554_01_C11 | 14,4 | 1 | 1,47 | 97,2 | 2,8 |
| 2554_01_D10 | 13,2 | 1 | 1,47 | 96,8 | 3,2 |
| 2554_01_E05 | 9,8 | 2 | 1,46 | 96,9 | 3,1 |
| 2555_02_D09 | 14,8 | 2 | 1,46 | 97,0 | 3,0 |
| 2555_02_D02 | 7,3 | 1 | 1,49 | 100,0 | 0,0 |
| 2555_01_B09 | 16,5 | 1 | 1,47 | 96,0 | 4,0 |
| 2554_01_D05 | 13,9 | 2 | 1,47 | 97,1 | 2,9 |
| 2554_01_C07 | 14,0 | 2 | 1,47 | 96,6 | 3,4 |
| 2558_01_E08 | 13,1 | 1 | 1,47 | 96,7 | 3,3 |
| 2554_02_G10 | 13,6 | 1 | 1,47 | 96,9 | 3,1 |
| 2558_02_F12 | 12,9 | 1 | 1,47 | 96,6 | 3,4 |
| 2555_02_D05 | 11,4 | 2 | 1,47 | 100,0 | 0,0 |
| 2551_02_E01 | 11,3 | 2 | 1,52 | 95,2 | 4,8 |
| 2558_02_H05 | 10,0 | 2 | 1,49 | 96,6 | 3,4 |
| 2558_02_G01 | 8,0 | 3 | 1,48 | 100,0 | 0,0 |
| 2558_02_G10 | 17,2 | 2 | 1,48 | 96,4 | 3,6 |
| 2551_01_B01 | 12,2 | 1 | 1,43 | 96,5 | 3,5 |
| 2554_02_H01 | 9,1 | 1 | 1,48 | 96,5 | 3,5 |
| 2554_02_G12 | 13,9 | 2 | 1,56 | 100,0 | 0,0 |
| 2554_01_D01 | 9,9 | 2 | 1,51 | 97,1 | 2,9 |
| 2551_01_A01 | 2,7 | 7 | 1,49 | 100,0 | 0,0 |
| 2554_02_G07 | 15,9 | 1 | 1,48 | 96,6 | 3,4 |
| 2554_02_F12 | 11,4 | 1 | 1,47 | 95,3 | 4,7 |
| 2555_01_B01 | 19,7 | 1 | 1,48 | 95,4 | 4,6 |
| 2558_01_E04 | 12,1 | 1 | 1,51 | 95,6 | 4,4 |
| 2558_02_H06 | 9,3 | 1 | 1,47 | 100,0 | 0,0 |
| 2558_01_E03 | 20,1 | 1 | 1,47 | 96,1 | 3,9 |
| 2558_01_D11 | 15,4 | 1 | 1,47 | 97,2 | 2,8 |
| 2558_02_G07 | 17,9 | 0 | 1,46 | 96,4 | 3,6 |
| 2555_01_B04 | 17,3 | 1 | 1,48 | 97,4 | 2,6 |
| 2558_02_G02 | 20,1 | -1 | 1,48 | 96,2 | 3,8 |
| 2558_01_E02 | 10,7 | 2 | 1,48 | 100,0 | 0,0 |
| 2555_02_D06 | 21,4 | 0 | 1,46 | 96,9 | 3,1 |
| 2554_01_D09 | 7,3 | 2 | 1,46 | 100,0 | 0,0 |
| 2555_02_D07 | 17,2 | 0 | 1,48 | 96,2 | 3,8 |
| 2558_01_D12 | 15,6 | 1 | 1,47 | 95,9 | 4,1 |
| 2558_02_G03 | 14,4 | 1 | 1,49 | 91,1 | 8,9 |
| 2558_01_E10 | 16,7 | 0 | 1,48 | 96,6 | 3,4 |
| 2558_02_G04 | 17,6 | 0 | 1,46 | 97,3 | 2,7 |
| 2554_01_D04 | 3,8 | 3 | 1,47 | 100,0 | 0,0 |
| 2554_02_F04 | 18,9 | - | 1,49 | 95,5 | 4,5 |
| 2555_02_C08 | 16,2 | - | 1,55 | 97,2 | 2,8 |
| 2558_02_G05 | 15,1 | - | 1,47 | 97,0 | 3,0 |
| 2555_01_A08 | 12,3 | - | 1,48 | 96,1 | 3,9 |
| 2558_02_F10 | 6,9 | - | 1,47 | 100,0 | 0,0 |
| 368_01_C05 | 14,7 | 0 | 1,45 | 97,5 | 2,5 |

**Fig. S1.**
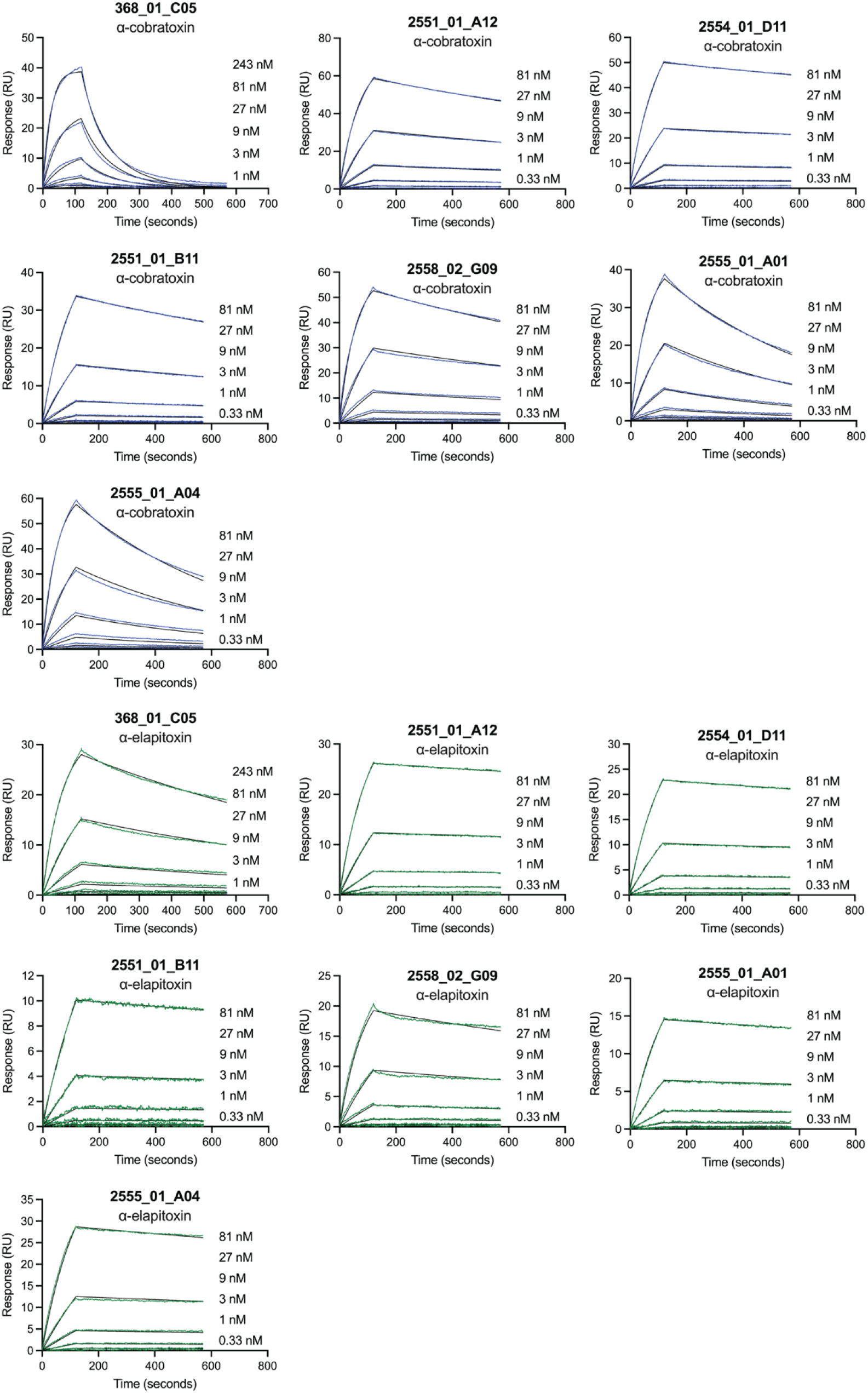
Affinity measurements using surface plasmon resonance. Sensograms illustrating affinity measurements of the top six affinity matured antibodies as well as the parent on α-cobratoxin and α-elapitoxin immobilized on a CM5 sensor.

**Fig. S2.**
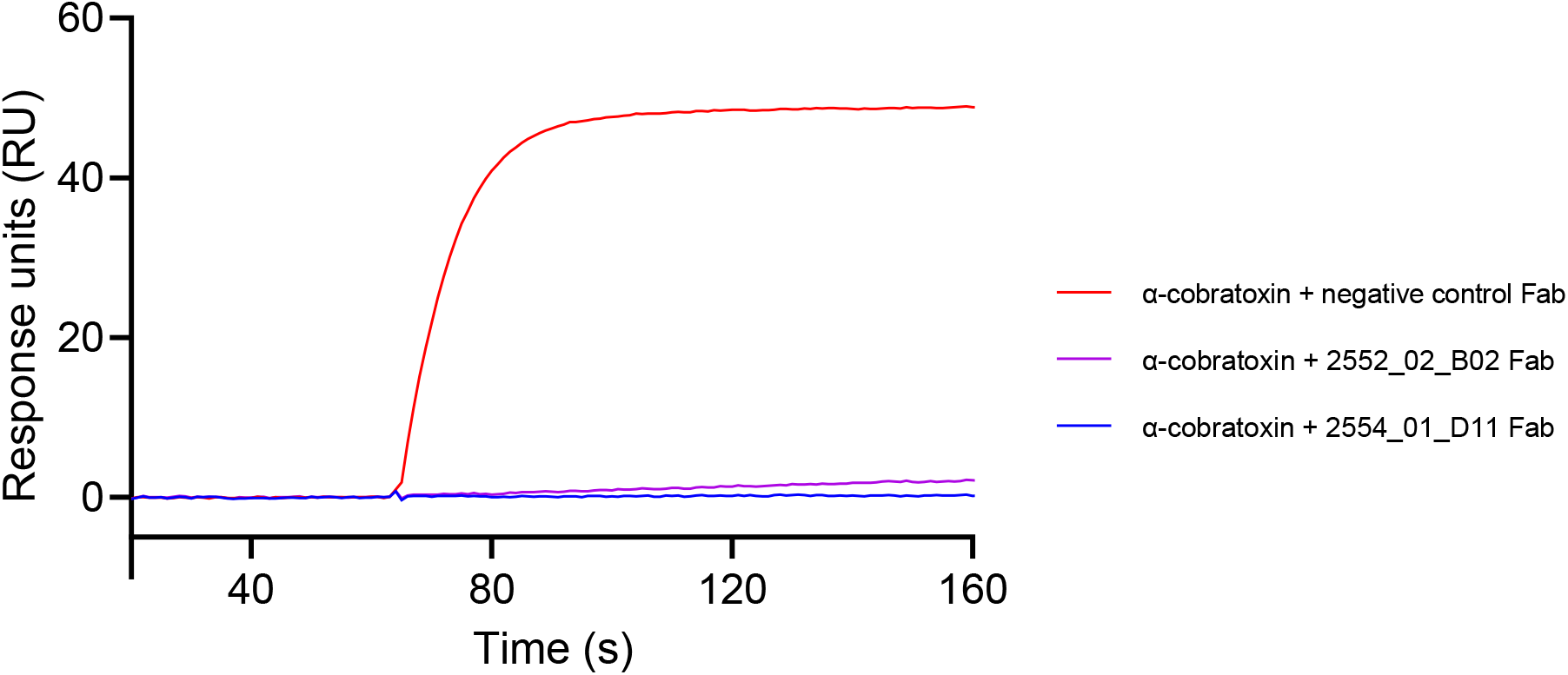
Epitope binning of 2554_01_D11 and 2552_02_B02 anti-α-cobratoxin Fabs by SPR. Sensorgrams of α-cobratoxin interacting with immobilized 2554_01_D11 Fab in the presence of 2552_02_B02 Fab. 200 nM of Fab was pre-incubated for 30 min with 20 nM α-cobratoxin before being flowed over immobilized 2554_01_D11 for 120s. A non α-cobratoxin specific Fab and the 2554_01_D11 Fab were included as negative and positive controls, respectively.

**Fig S3.**
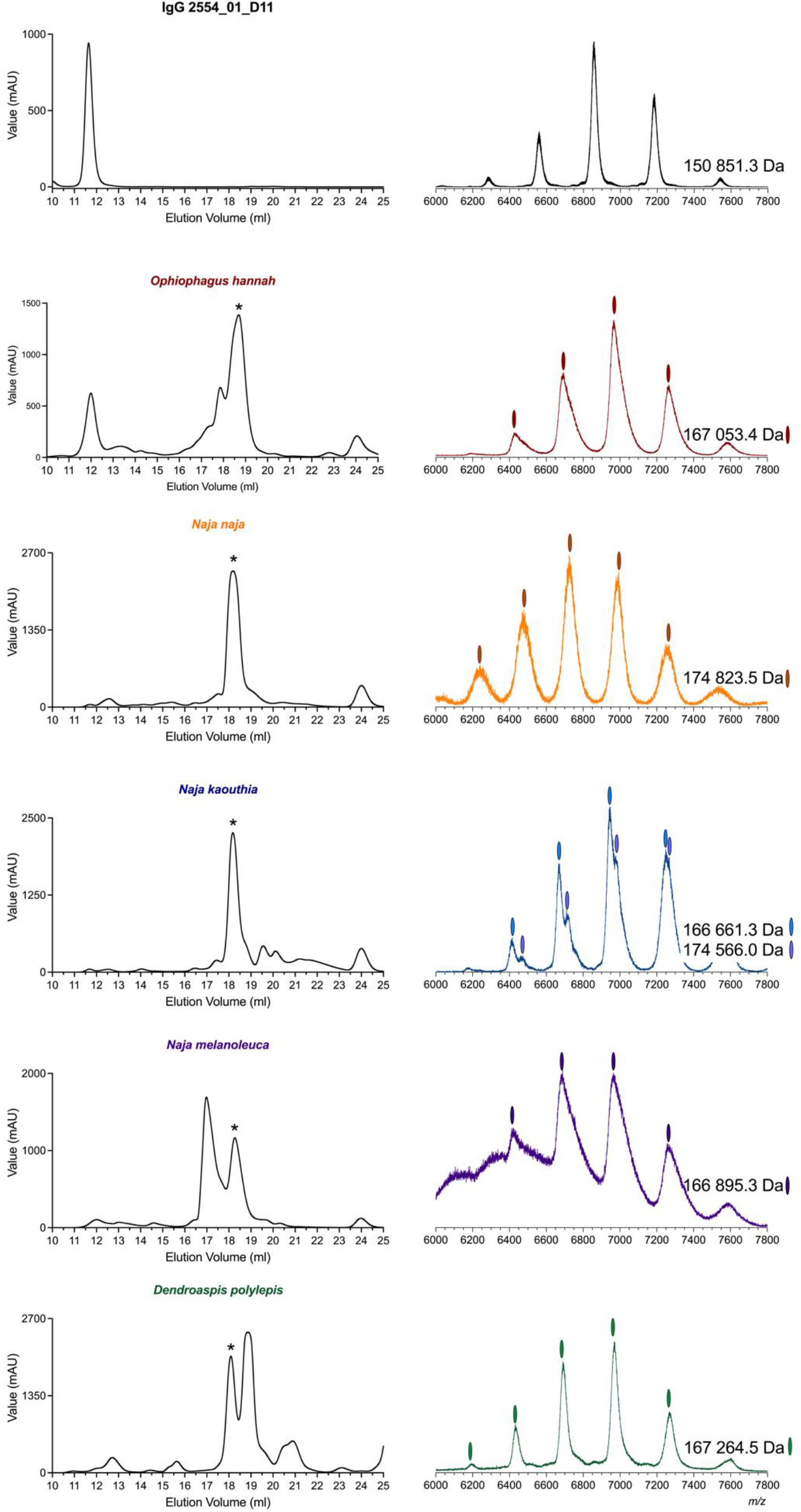
Size exclusion chromatograms of the snake whole venoms and native mass spectra of toxin:antibody complexes. Size exclusion chromatograms of IgG 2554_01_D11 and five featured venoms accompanied by native mass spectra of IgG 2554_01_D11 – toxin fraction with asterix from each SEC run.

